## Supporting_information for "Visualization of liquid-liquid phase transitions using a tiny G-quadruplex binding protein"

### Supporting methods

***In silico* RNA motif extraction**: Library-1 from the previously published paper[1] was used for all experiments. All motifs including human pre-miRNA in the library were extracted from miRBase. As a control, RNAG4 motifs were included in the library.

***Oligo template pool and DNA barcode microarray design***: To generate the RNA structure library, single-stranded DNA sequences were used as templates for RNA probes, including each RNA structure extracted from the datasets. The templates were synthesized by OLIGONUCLEOTIDE LIBRARY SYNTHESIS (OLS, Agilent Technologies). Each template DNA included five components arranged in the following order from the 3′ end: CC + T7 promoter sequence, Barcode sequence, Stabilizing stem structure sequence Forward, RNA structure region, and Stabilizing stem structure sequence Reverse as previously described1. The size of the oligo templates was limited to 170nt for the RNA structure library. After assigning barcodes to the RNA structure, we converted the barcodes to antisense DNA strands. The resulting DNA strands were submitted to the SureDesign server as CGH custom array design services (Agilent Technologies). We set the Probe Replication Factor to 5× for the RNA structure library. Then, a custom CGH DNA microarray was purchased in the 8× 60 K array format.

***In vitro* transcription of the RNA structure library**: To produce the RNA structure library, the MEGAshortscript T7 Transcription Kit (Thermo Fisher Scientific) was used according to the manufacturer's instructions. The reaction solution was gently mixed and had a total volume of 20 μL, containing ssDNA templates and the ssDNA coding T7 promoter sequence. The contents were mixed thoroughly by gently flicking the tube, and the reaction mixture was briefly microfuged to collect it at the bottom of the tube. The reaction was incubated at 37°C for 20 hours using the ProFlex PCR System (Thermo Fisher Scientific) with a lid heated to 105°C to prevent evaporation. Following incubation, 2 μL of TURBO DNase (Thermo Fisher Scientific) was added to the reaction solution, mixed by pipetting, and incubated at 37°C for 15 minutes. The RNA products were then purified using the Zymo RNA Clean and Concentrator kit (Zymo Research).

***3’ - terminal Cy5 labeling***: To detect and quantify RNA probes on a microarray, the library of RNA probes was labeled with a fluorescent dye at the 3' end. A reaction mixture containing 1× T4 Ligase Buffer (Thermo Fisher Scientific), 100 μM pCp-Cy5 (Jena Bioscience), 10 μM RNA structure library, and 0.5 U/μL T4 RNA Ligase (Thermo Fisher Scientific) was prepared and incubated at 16°C for 48 hours under light-shielded conditions. The labeled RNA probes in the library were then purified using the Zymo RNA Clean and Concentrator kit (Zymo Research). The labeled RNA structure library was stored at -28°C until further use.

***Hybridization and microarray scanning***: To prepare the RNA samples for hybridization, 18 μL of enriched RNA was mixed with 4.5 μL of 10× Blocking Agent (Agilent Technologies) and 22.5 μL Hi-RPM Hybridization Buffer (Agilent Technologies) by vortexing and centrifuging to collect the reaction mixture at the bottom of the tube. The samples were then incubated for 5 minutes in a heat block set at 104 °C and immersed in ice water for an additional 5 minutes. The RNA samples were applied to an 8×60 K Agilent microarray gasket slide (Agilent Technologies), which was assembled with a CGH custom array 8×60 K (Agilent Technologies) using SureHyb. Hybridization was performed in a hybridization oven (Robbins Scientific) for 20 hours at 55.5 °C while rotating at 20 rpm. After hybridization, the microarray slide was washed for 5 minutes with Gene Expression Wash Buffer 1 (Agilent Technologies) in a glass container at room temperature. The microarray slide was then transferred to a glass container containing Gene Expression Wash Buffer 2 (Agilent Technologies) and immersed in a thermostat bath at 37 °C for another 5 minutes. The fluorescence image data of the microarray were obtained using SureScan (Agilent Technologies). Finally, the captured images of the microarray slide were converted to numeric fluorescence intensities of each spot using Feature Extraction (Agilent Technologies) and GeneSpringGX (Agilent Technologies).

### Supplementary figures

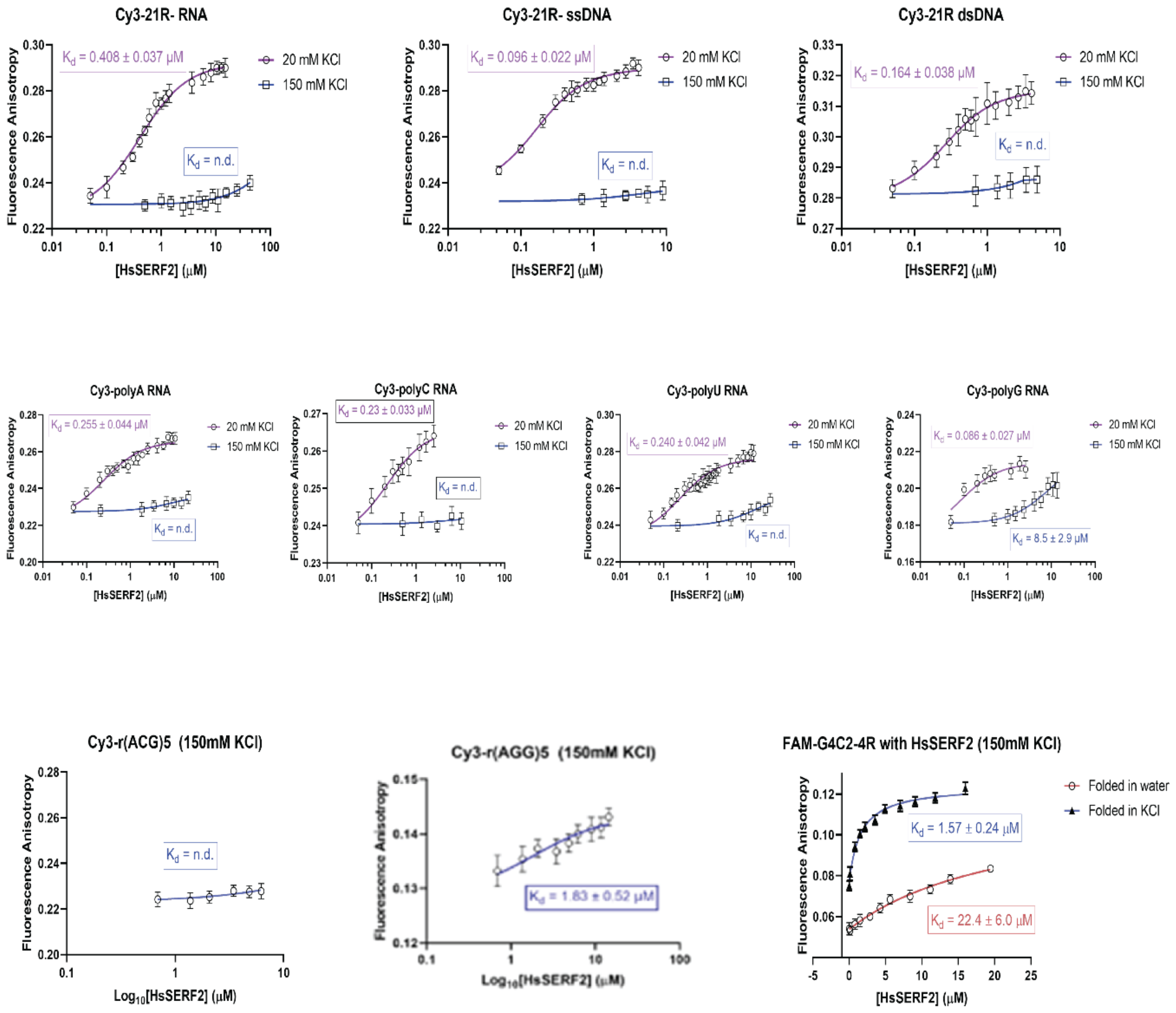


**Figure S1. SERF2 binds specifically to folded rG4s.**

Fluorescence anisotropy titration of human SERF2 protein into Cy3 labeled 21R- single-stranded RNA, single-stranded DNA, and double-stranded DNA; Cy3 labeled poly-tract RNAs with 21 nucleotides in length; Cy3 labeled (ACG)5 and (AGG)5 RNA, and 6-FAM labeled 4-repeat G4C2. SERF2 presented a salt-dependent and low micromolar binding affinity to different DNA and RNA oligonucleotides, with a relatively higher affinity at low salt concentration. Data are presented as mean values ± standard deviations.

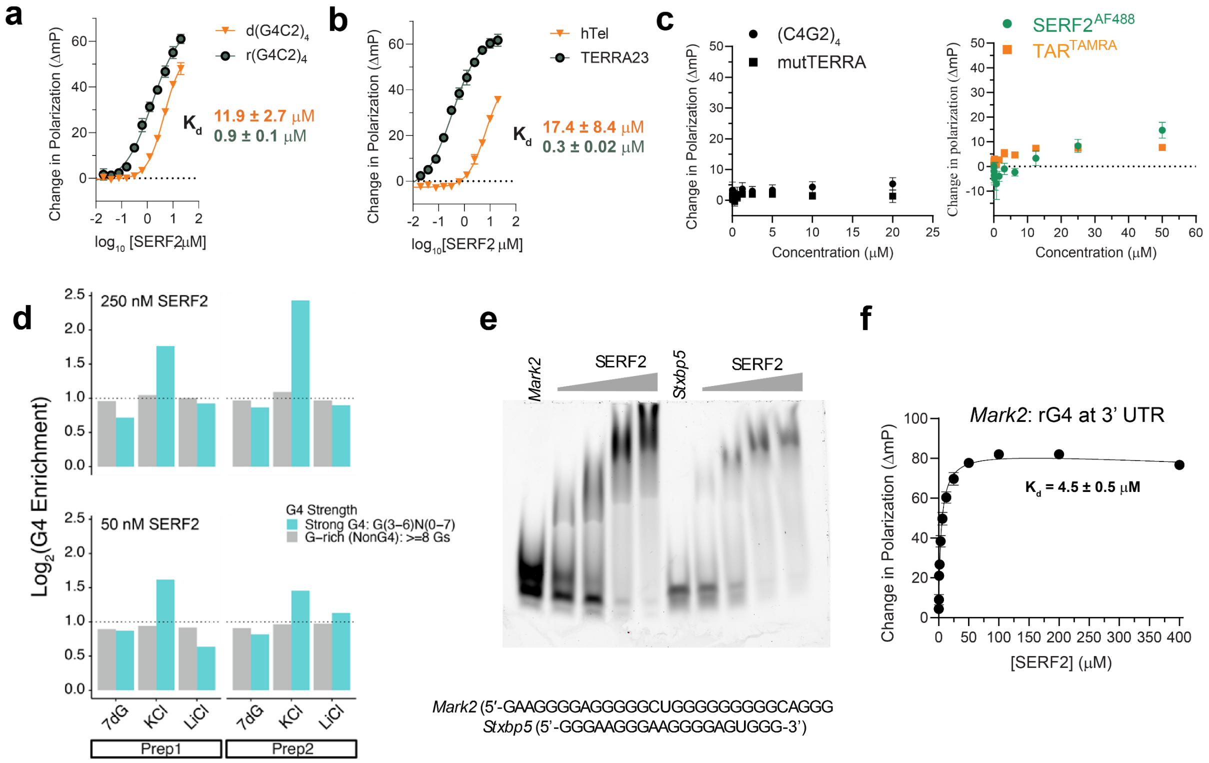


**Figure S2. Fluorescence polarization assay comparing the binding affinity of SERF2 to DNA and RNA G4s. a-b** Fluorescence polarization (FP) binding curves of SERF2 with 4-repeat DNA compared to its binding to RNA G4C2 (**a**), and the 4-repeat human telomeric repeat sequence TTAGGG compared to the TERRA23 RNA (**b**). **c** SERF2 showed very weak binding to non rG4 forming structured TAR (right) or unstructured (CCCCGG)4 and mutated TERRA (UUACCG)4 (left) repeat RNA sequences. FP assays were done using SERF2-AF488 for all RNA sequences including TAR. Additional FP test using TAR-TAMRA confirms SERF2 binds weakly to TAR. All fluorescence polarization assays (except TAR-TAMRA) were done with an increasing concentration of SERF2 against a fixed 100 nM concentration of DNA or RNA G4s dissolved in 20 mM NaPi, 100 mM KCl at room temperature. The binding constants were calculated by fitting the fluorescence polarization curves using a binding saturation one-site total model in GraphPad Prism. **d** Enrichment of rG4sin RNA bind-n-seq analysis using two different preparations of RNA pools prepared in KCl, or LiCl solutions, or synthesized using 7-deaza. RNA sequences containing 3 or more guanines in the G-tetrad are referred to as strong G4s and sequences with ≥8 guanines but lacking a defined G4 forming motif, as defined by QGRS mapper program, are referred to as non rG4s. An enrichment analysis done for two different SERF2 concentrations (250 nM, top; 50 nM, bottom) shows a higher rG4 enrichment for the highest (250 nM) SERF2 concentration in the KCl solution. **e** The EMSA gel-shift assay shows binding of SERF2 (5, 10, 20 and 40 μM) to rG4s (10 μM) derived from the 3’ UTR regions of *Mark2* and *Stxbp5* genes. **f** Binding affinity measurement (FP assay) between SERF2 and 100 nM 6-FAM labeled rG4 present at 3’ UTR in *Mark2* gene. FP assay data shown in **a-c** and **f** are presented as mean values ± standard deviations obtained from three independent replicates.

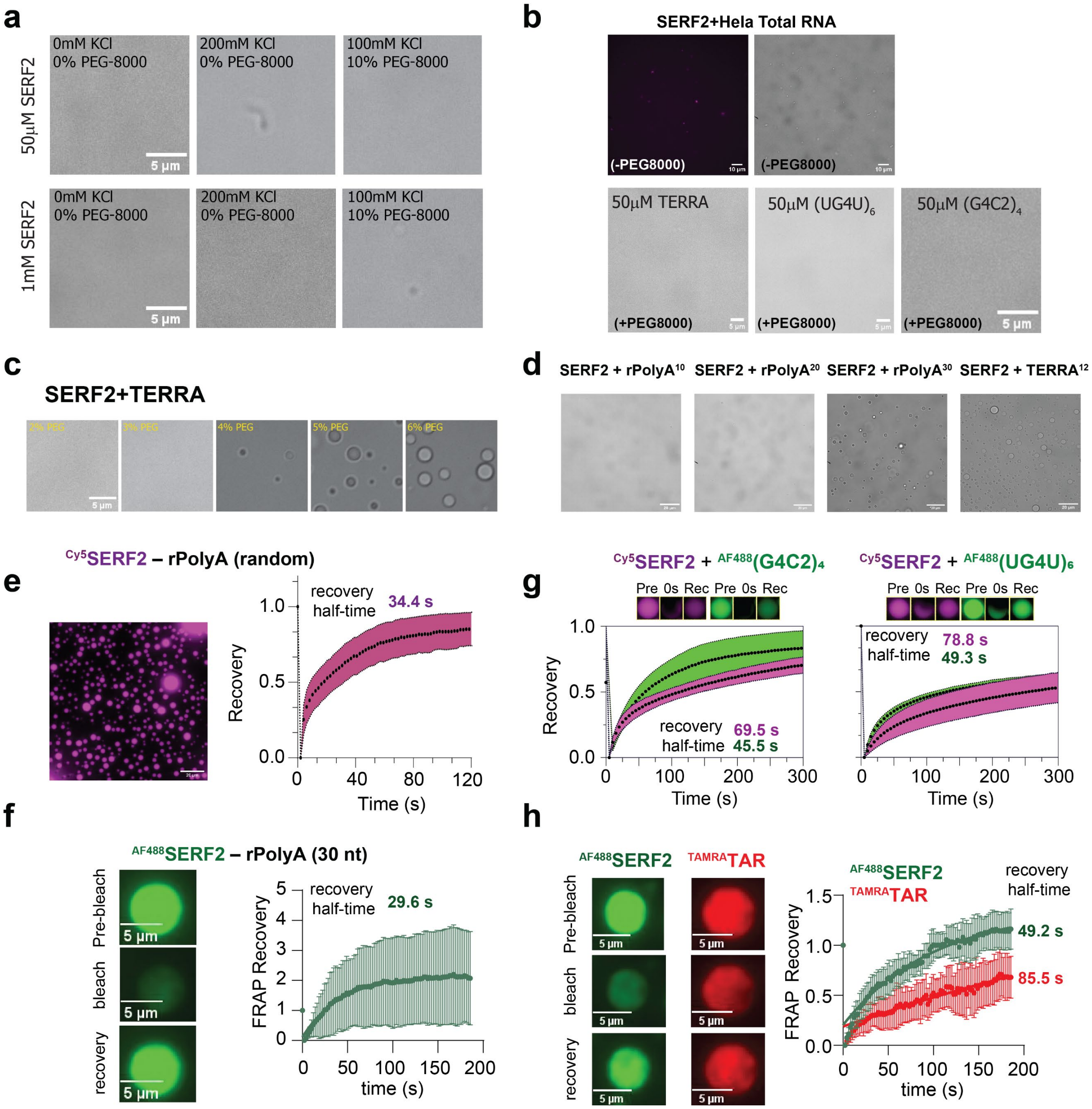
a
50μM SERF2
1mM SERF2
0mM KCl
0% PEG-8000
200mM KCl
0% PEG-8000
100mM KCl
10% PEG-8000
5 μm
b
SERF2+Hela Total RNA
(-PEG8000)
50μM TERRA
50μM (UG4U)6
50μM (G4C2)4
(+PEG8000)
5 μm
c
SERF2+TERRA
2% PEG
3% PEG
4% PEG
5% PEG
6% PEG
5 μm
d
SERF2 + rPolyA10
SERF2 + rPolyA20
SERF2 + rPolyA30
SERF2 + TERRA12
e
Cy5SERF2 – rPolyA (random)
recovery half-time 34.4 s
Recovery
Time (s)
g
Cy5SERF2 + AF488(G4C2)4
Cy5SERF2 + AF488(UG4U)6
Pre 0s Rec Pre 0s Rec
recovery half-time 69.5 s 45.5 s
recovery half-time 78.8 s 49.3 s
Recovery
Time (s)
f
AF488SERF2 – rPolyA (30 nt)
Pre-bleach
bleach
recovery
5 μm
recovery half-time 29.6 s
FRAP Recovery
time (s)
h
AF488SERF2
TAMRATAR
Pre-bleach
bleach
recovery
5 μm
recovery half-time
49.2 s
85.5 s
FRAP Recovery
time (s)

**Figure S3. Phase transition properties of SERF2. a** DIC images show no droplet formation for 50 µM and 1 mM SERF2 dissolved in 20 mM NaPi, pH 7.4 containing different salts and PEG8000 as indicated. **b** Images show all three RNA G4s do not form droplets on their own in 20 mM NaPi, 100 mM KCl, pH 7.4 buffer containing 10% PEG8000. **c** DIC images of 50 µM TERRA12 mixed with equimolar SERF2 dissolved in 20 mM NaPi, 100 mM KCl, pH 7.4 containing variable PEG8000 concentrations as indicated. **d** Liquid-liquid phase separation in 50 µM SERF2 mixed with 50 µM of 10-nucleotide polyA (left) or TERRA12 (right) in NaPi buffer containing 10% PEG8000 and 100 mM KCl. **e, f** Phase transition of 50 µM SERF2 mixed with random length polyA (**e**) RNA (700-3500 kDa) or 30-nucleotide polyA (**f**) dissolved in 20 mM NaPi, 100 mM KCl, pH 7.4 buffer containing 10% PEG8000 and 1:200 diluted Cy5- or AF488-SERF2 as indicated to provide the purple or green signal, respectively. The images on the left and the plot on the right show the FRAP recovery of SERF2 as a function of time. Standard deviations were calculated by analyzing 8 (**e**) and 4 isolated droplets (**f**) subjected to FRAP. **g** Two-component FRAP analysis was done to measure the recovery rates of 50 µM SERF2 mixed with equimolar concentrations of (G4C2)4 or (UG4U)6 rG4s in the SERF2-rG4s droplets (see Figure 2b). Standard deviations were calculated by analyzing 8 isolated droplets subjected to FRAP. The sample mixture is doped with 1:200 diluted Cy-5 labeled SERF2 (purple) and 6-FAM rG4 for fluorescence imaging (green). On the top of each FRAP plot, the pre-bleach, post-bleach (0 s), and recovered droplets (300 s) are shown. **h** FRAP recovery images and plots are shown for SERF2 binding to non rG4 structured HIV-1 TAR RNA. Standard deviations were calculated by analyzing 4 isolated droplets subjected to FRAP. All FRAP data shown in Fig. S3 were fitted in GraphPad Prism, using a non-linear regression, one-phase association model, to obtain the recovery halftime.

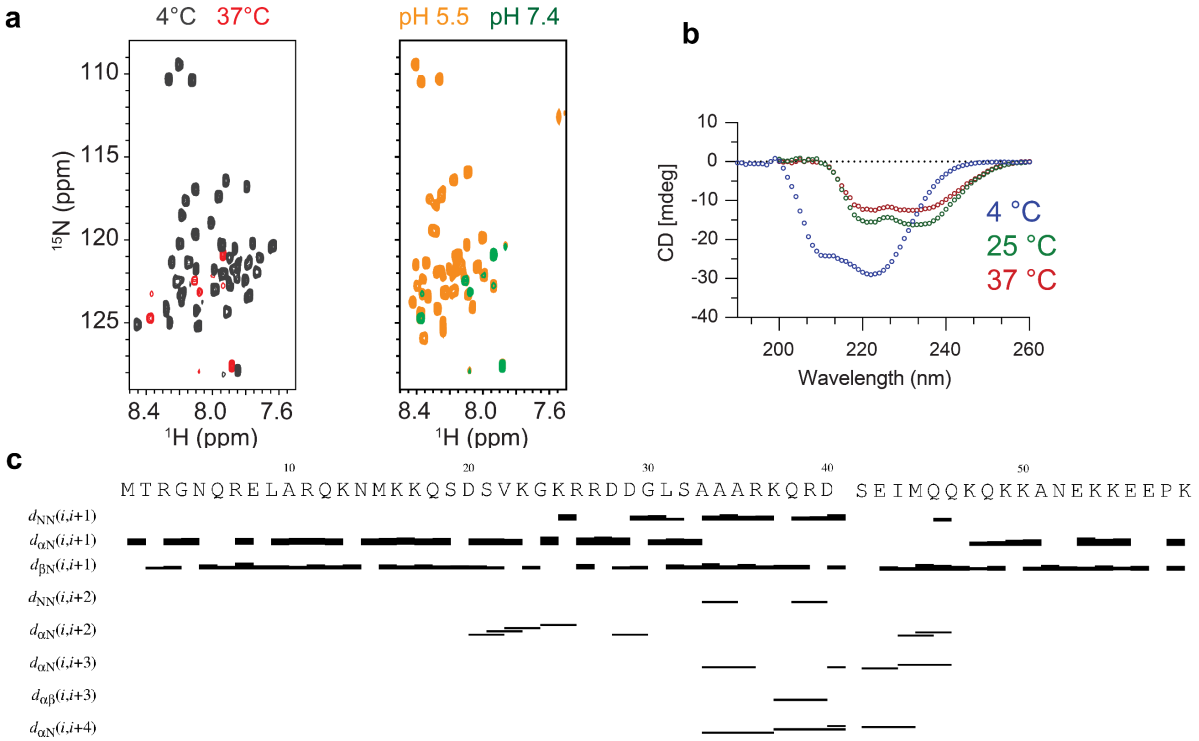


**Figure S4.** **SERF2 is partially disordered. a** Overlaid 1H-15N HSQC NMR spectra of 100 µM SERF2 recorded at 4 °C (black) or 37 °C (red and green) dissolved in 20 mM NaPi (pH 7.4), 100 mM KCl. The missing SERF2 peaks are detectable at 37 °C when dissolved in 20 mM d3-NaAc (pH 5.5),100 mM KCl (orange). The NMR data were collected on a Bruker 800 MHz in a buffer containing 8% $D_2O$. **b** Secondary structure analysis of human SERF2 by circular dichroism spectroscopy at the indicated temperature in 20 mM NaPi (pH 7.4), 100 mM KCl. **c.** NOE assignment plot generated by CYANA for SERF2, showing the positions of assigned NOEs along the amino acid sequence.

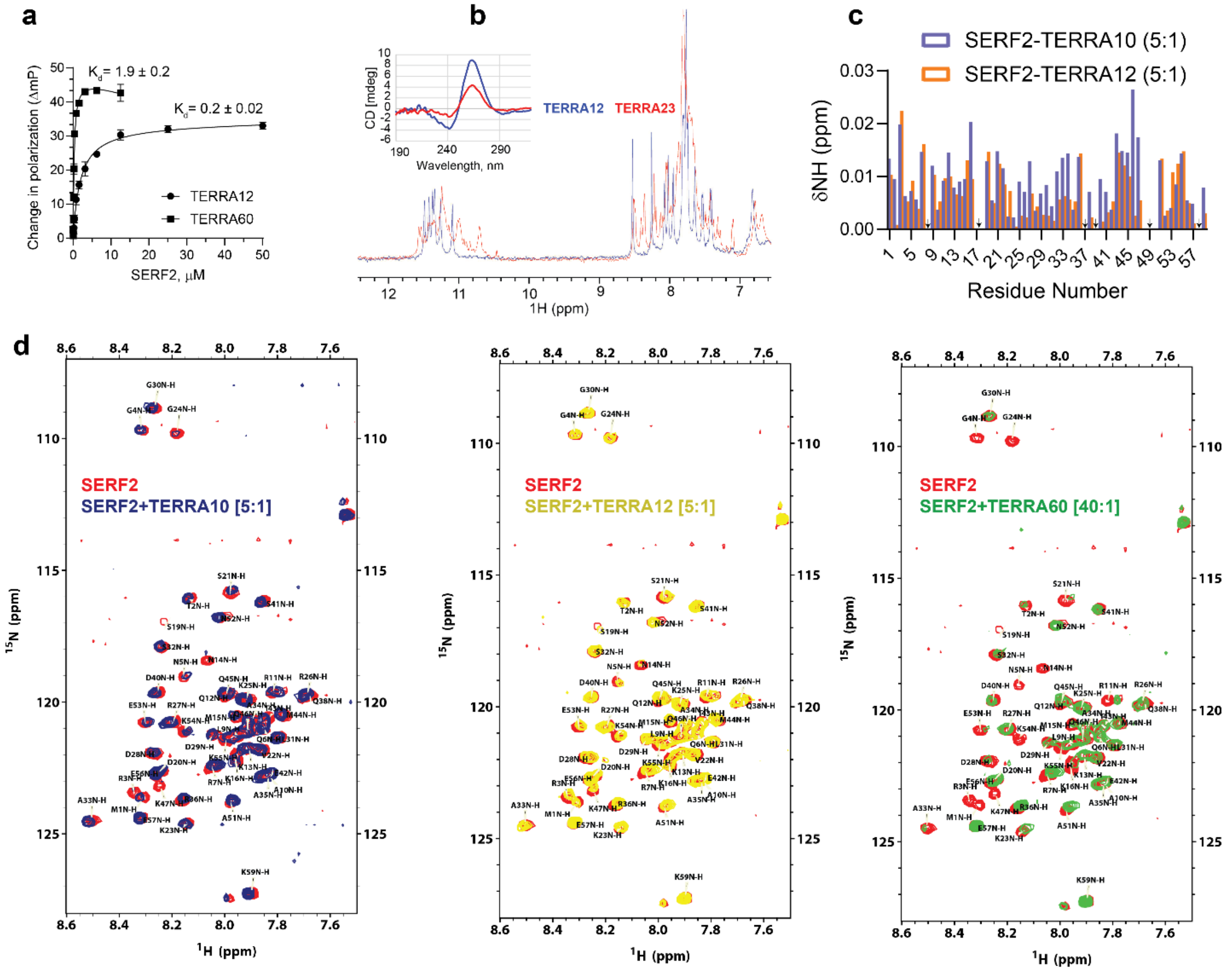


**Figure S5. SERF2 maintains a conserved binding interface across TERRA rG4s of varying lengths.** **a** Fluorescence polarization assay measured binding dissociation constant ($K_D$) using 200 nM SERF2-AF488 mixed with an increasing concentration of 12- or 60-nucleotide long TERRA12 or TERRA60. Data are presented as mean values ± standard deviations obtained from three independent replicates. **b** Overlaid proton NMR spectrum of 100 µM TERRA12 and TERRA23 dissolved in 20 mM NaPi (pH 7.4), 100 mM KCl, showing well-resolved imino protons for TERRA12 as compared to TERRA23. The figure inset verifies a stronger ~263 nm CD signal for 10 µM TERRA12 as compared to TERRA23, suggesting its relatively better parallel folding structure. **c** Comparison of 1H-15N chemical shift perturbations in SERF2 (50 µM) upon binding to TERRA10 and TERRA12 rG4s at a 5:1 RNA:protein ratio (empty regions in the graph marked with an arrow correspond to unassigned peaks). Chemical shift perturbation profiles display similar trends across the SERF2 sequence, indicating a conserved binding interface is involved in TERRA10, 12 or 23 (see Figure 3b) rG4 binding. The 1H-15N HSQC spectra are shown in "**d**". **d** Overlaid 1H-15N HSQC NMR spectra of 50 µM SERF2 mixed with different lengths of TERRA RNA, as indicated, recorded at 4 °C. All NMR samples were prepared in 20 mM NaPi (pH 7.4), 100 mM KCl.

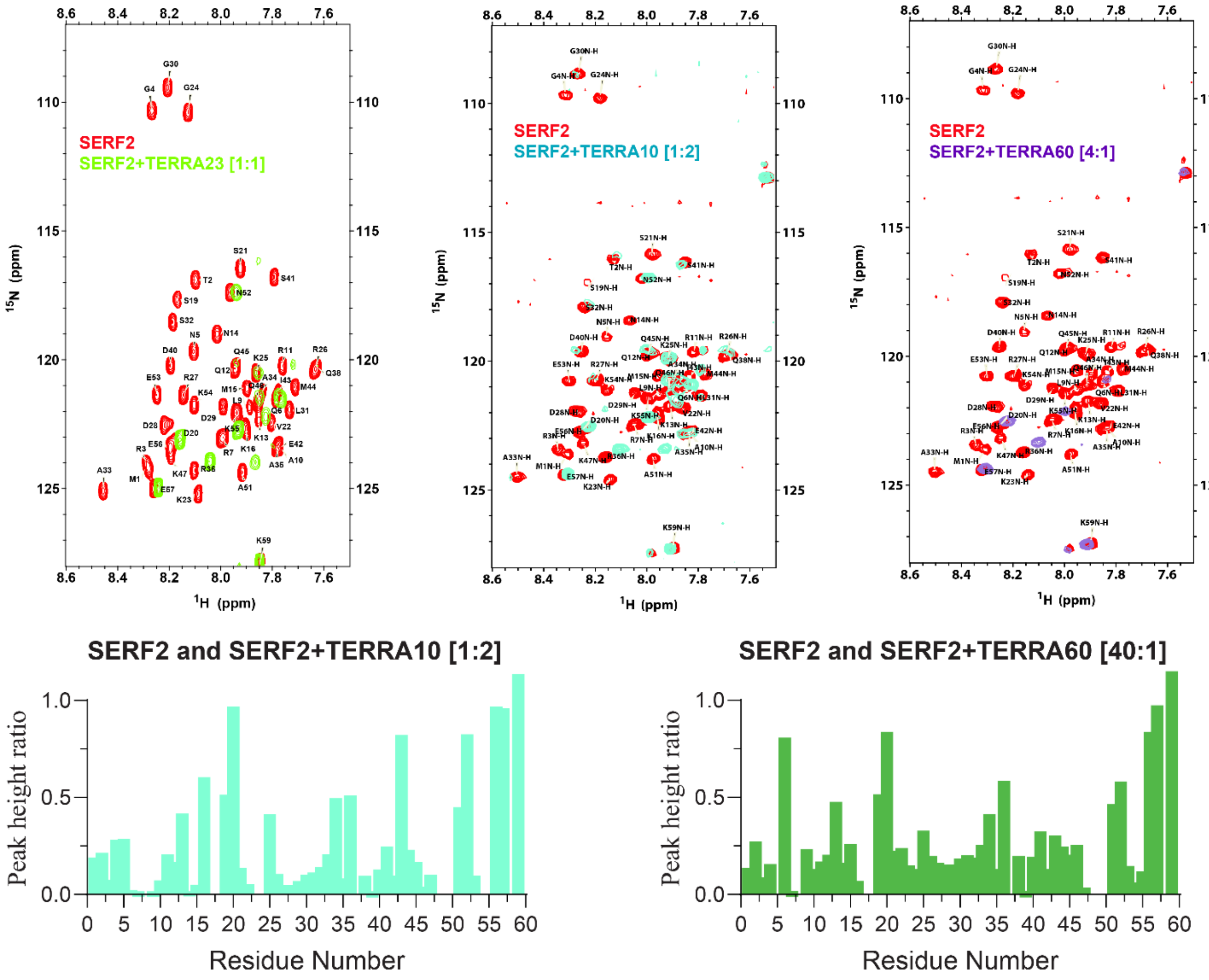


**Figure S6. Signal broadening in 1H-15N HSQC spectra reveals binding of SERF2 to TERRA rG4s of varying lengths.** Overlay of 1H-15N HSQC spectra of SERF2 upon titration with 10-, 23-, and 60-nucleotide TERRA rG4 at high RNA concentrations as indicated. All spectra exhibit progressive signal broadening of SERF2 cross-peaks, indicative of reduced tumbling, possibly due to RNA binding and increased complex molecular weight. The bar graphs show the peak height ratio of SERF2 in the presence of TERRA10 and TERRA60 compared to the free protein. A reduction in the peak height ratio for many residues indicates significant signal broadening in the 1H-15N HSQC spectra. This broadening is indicative of a slower molecular tumbling rate, which is consistent with the formation of a larger protein-RNA complex upon binding. The left plot corresponds to a 1:2 protein:TERRA10 ratio (spectra shown in the upper middle panel), while the right plot corresponds to a 40:1 protein:TERRA60 ratio (corresponding spectra are shown in Figure S5d, bottom right panel).

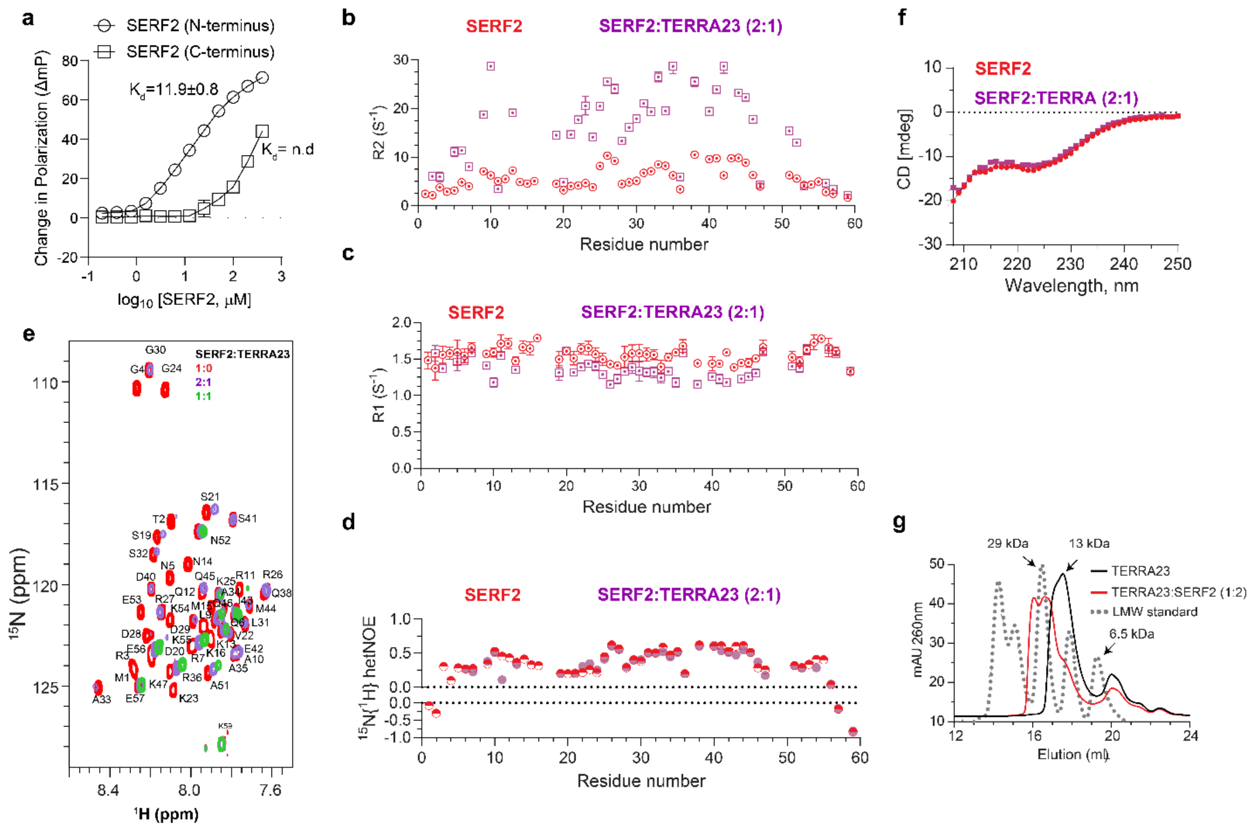


**Figure S7. SERF2 N-terminal domain mediates rG4 binding without inducing major structural rearrangements but increasing structural rigidity. a** Fluorescence polarization assay measuring the binding dissociation constant ($K_D$) of N-terminal (1-32) and C-terminal (31-59) SERF2 to 100 nM 6-FAM TERRA23 rG4. The fluorescence polarization binding data were generated from three independent replicates, and curves were fitted by GraphPad Prism non-linear regression curve fit and one-site fitting model analysis. Data are presented as mean values ± standard deviations. **b** 15N $R_2$ relaxation rates for SERF2 in the absence (red) and presence (purple) of TERRA23 rG4 RNA. A significant increase in $R_2$ is observed upon rG4 binding, indicating reduced molecular tumbling and potential complex formation. **c** 15N $R_1$ relaxation rates for SERF2 without (red) and with (purple) TERRA23 rG4. Minimal changes suggest no substantial alterations in fast timescale dynamics upon binding. **d** $\{^1H\}$-15N heteronuclear NOE values for SERF2 in the absence (red) and presence (purple) of TERRA23 rG4. The largely unchanged NOE values indicate that SERF2 retains flexibility in its disordered regions despite binding. Error bars represent the standard error from per-residue exponential fitting of peak intensities in NMRFAM-Sparky. **e** Overlaid 1H-15N HSQC NMR spectra of 100 μM SERF2 (red) mixed with 50 μM TERRA23 rG4 (purple) and 100 μM TERRA23 rG4 (green). The NMR data were collected at 4 °C on a Bruker 800 MHz. **f** Far-UV CD spectra of 40 μM SERF2 (205-250 nm) in the absence (red) and presence (purple) of 20 μM TERRA23 rG4 at 25 °C. Spectra were baseline-corrected by subtracting the TERRA23 rG4 signal. The overlay shows no significant secondary structure change in SERF2 upon rG4 binding. CD measurements were performed using a JASCO spectropolarimeter with 8 averaged scans. **g** Size-exclusion chromatography profiles of 20 μM TERRA23 rG4 mixed with 40 μM of SERF2 dissolved in 20 mM NaPi, 100 mM KCl, pH 7.4 buffer. The size exclusion chromatography profile shown in the dashed line corresponds to a low-molecular-weight (LMW) standard.

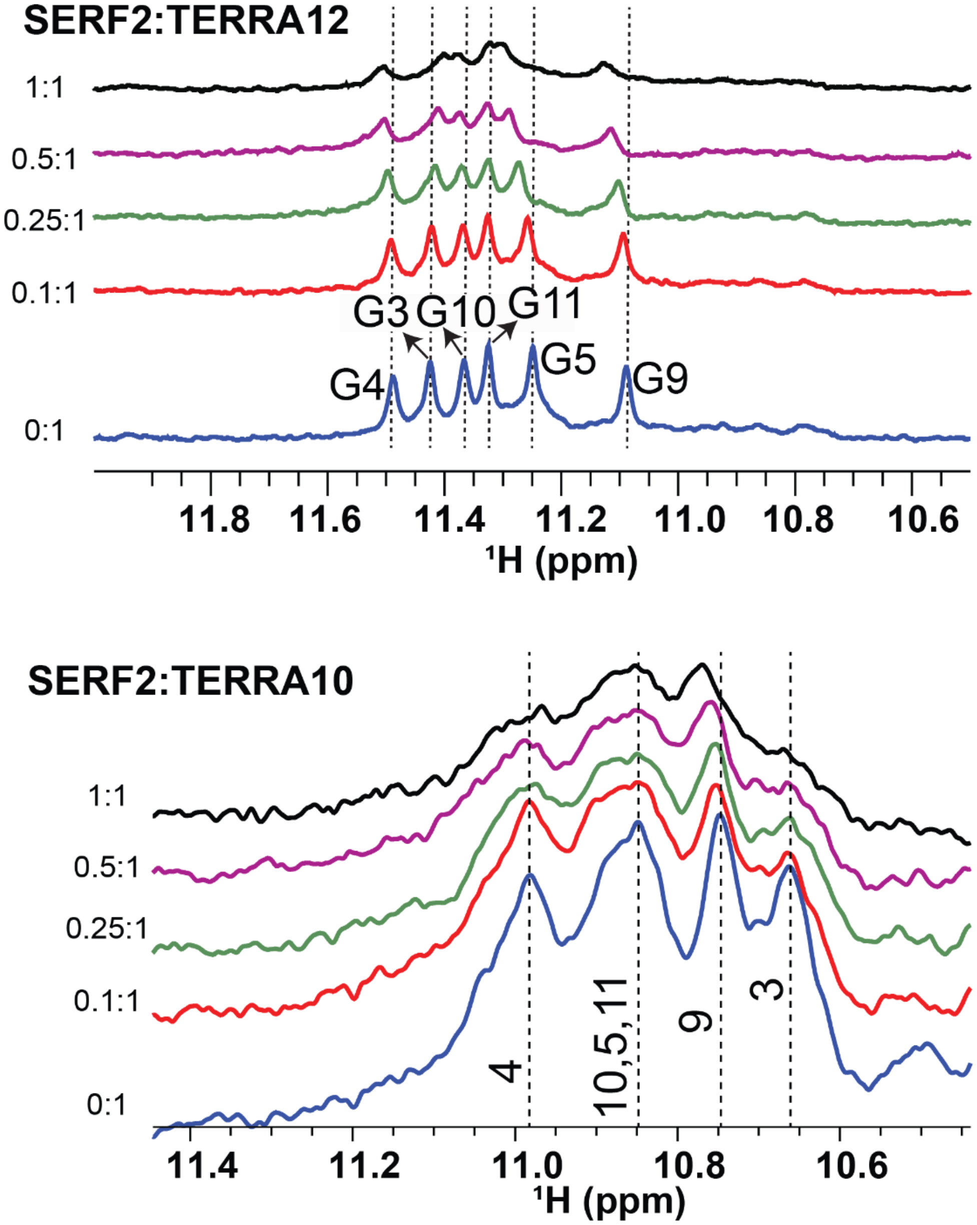


**Figure S8. 1D proton NMR titration measurements to map SERF2 binding sites on TERRA rG4.** 1D $^{1}H$ NMR titrations of TERRA12 (200 μM, top) or TERRA10 (100 μM, bottom) rG4 oligonucleotides with increasing SERF2 concentrations (0:1 to 1:1 molar ratios). Progressive chemical shift perturbations and line broadening are observed in the imino proton region (10.5-12.5 ppm), indicating SERF2 binding induced changes in Hoogsteen base-pairing. Dashed lines trace shifting resonances corresponding to specific guanine residues, confirming SERF2 binding to the G-quadruplex core.

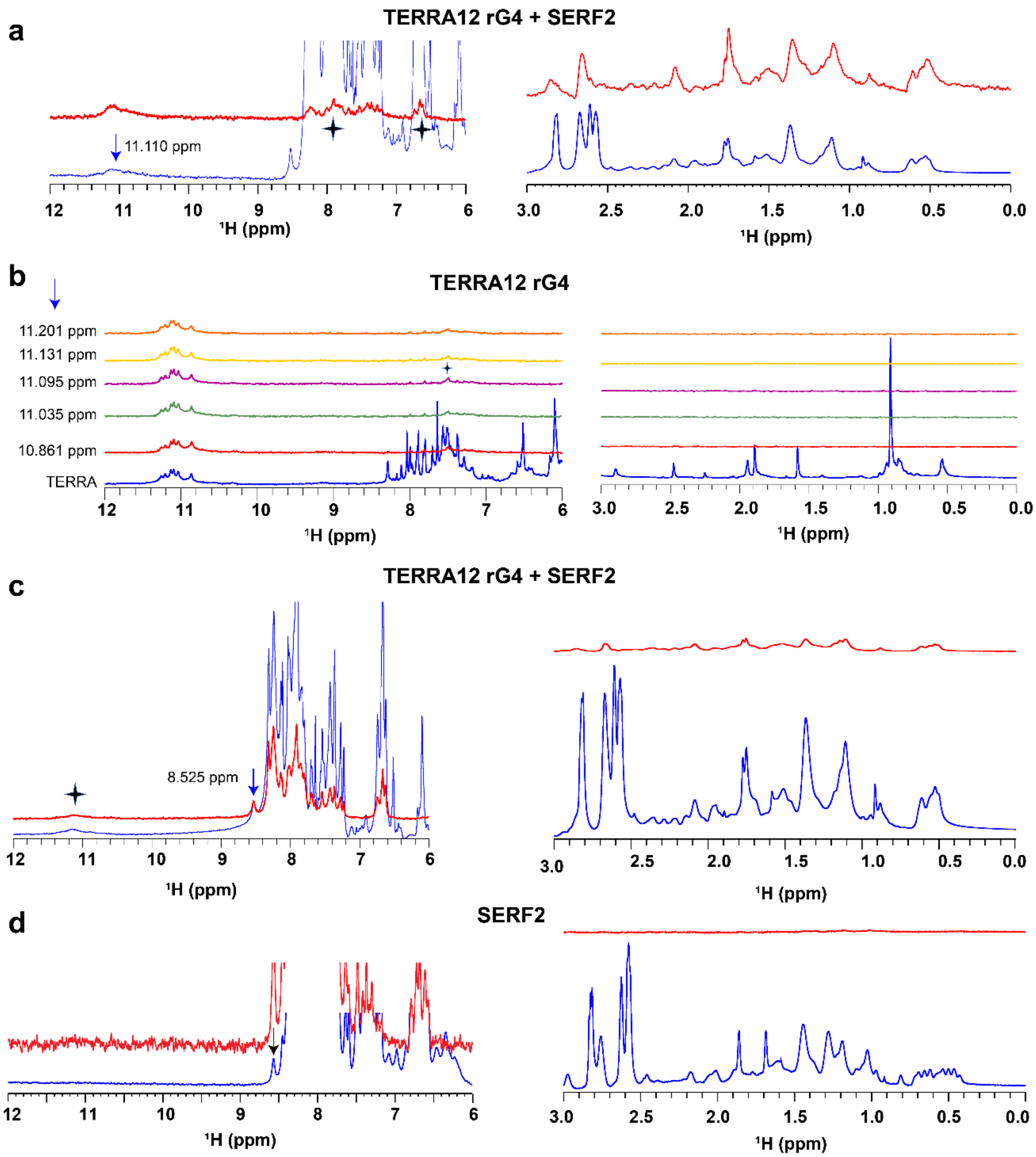


**Figure S9. SERF2 interacts with TERRA rG4 via specific binding contacts and shows direct proton-proton magnetization transfer in the complex. a** Saturation transfer difference NMR of SERF2-TERRA rG4 complexes shows magnetization transfer in the amino (~7-8.5 ppm, left) or aliphatic (0-3 ppm, right) proton regions upon saturating TERRA12 rG4 imino proton at ~11.11 ppm (blue arrow). The emergence of STD signals confirms close spatial proximity and binding between SERF2 and rG4s. The asterisks highlight selective transfer, consistent with intermolecular NOE contacts in the complex. The reference and STD difference NMR spectra are shown in blue and red, respectively. **b** STD NMR spectra of TERRA rG4 in isolation show very weak or no magnetization transfer upon saturating different imino proton signals as indicated (blue arrow) for amino (left) and aliphatic (right) regions. The reference spectrum is shown in blue, the difference spectra for distinct saturated imino protons are shown in different colors. **c** Saturation of the isolated amide proton resonance corresponding to A33 residue in SERF2 show magnetization transfer to TERRA12 rG4 imino regions (left) and aliphatic regions (right). The reference and STD difference NMR spectra are shown in blue and red, respectively. **d** Saturation of SERF2 in isolation at peak region ~8.5 ppm shows no magnetization transfer to TERRA rG4 imino regions or aliphatic regions (red spectrum). Arrows and asterisks respectively indicate saturated and transferred signals.

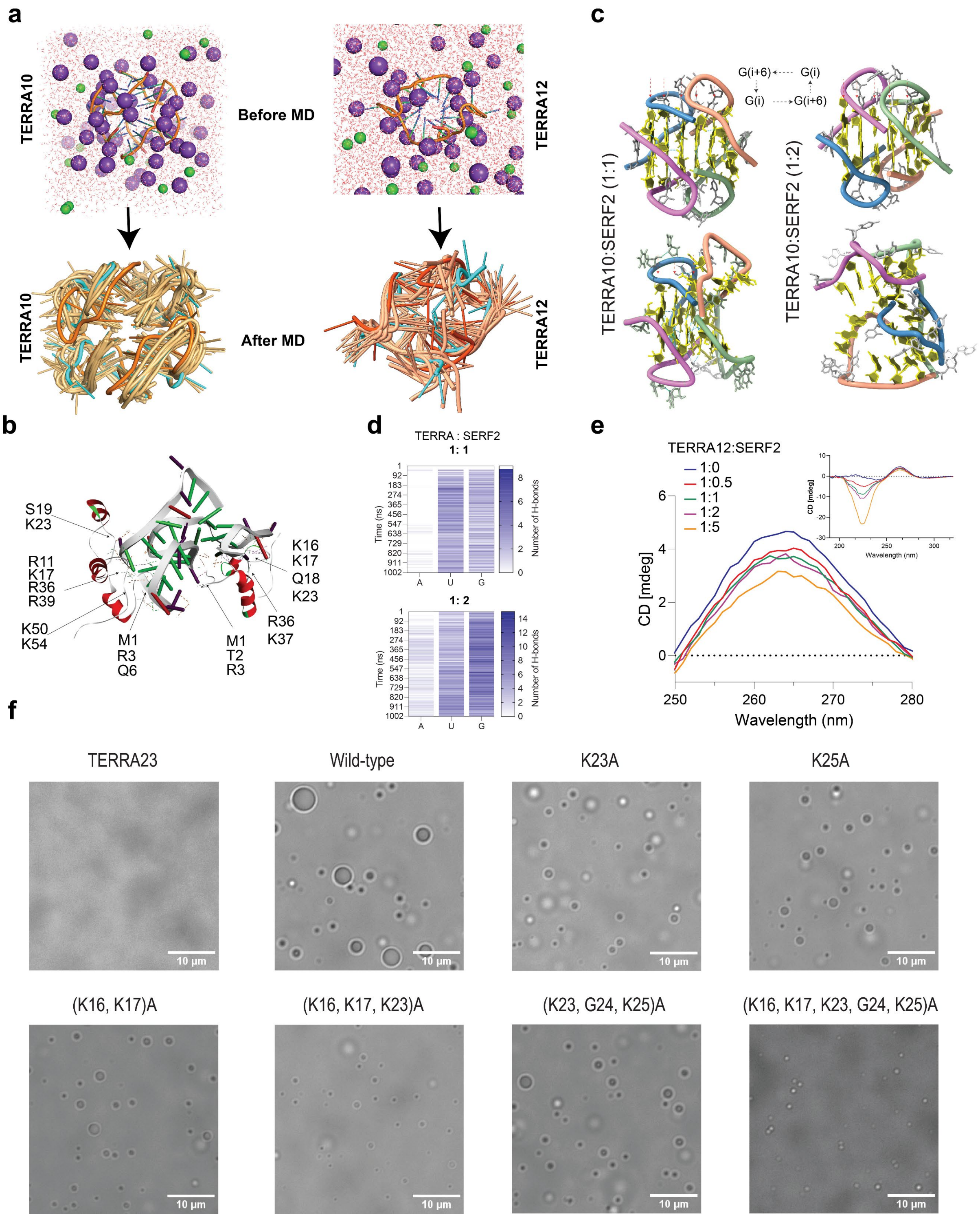
a
TERRA10
Before MD
TERRA12
TERRA10
After MD
TERRA12
c
G(i+6)
G(i)
G(i)
G(i+6)
TERRA10:SERF2 (1:1)
TERRA10:SERF2 (1:2)
b
S19
K23
R11
K17
R36
R39
K50
K54
M1
R3
Q6
M1
T2
R3
K16
K17
Q18
K23
R36
K37
d
TERRA : SERF2
1: 1
1: 2
Time (ns)
Number of H-bonds
A
U
G
e
TERRA12:SERF2
1:0
1:0.5
1:1
1:2
1:5
CD [mdeg]
Wavelength (nm)
f
TERRA23
Wild-type
K23A
K25A
(K16, K17)A
(K16, K17, K23)A
(K23, G24, K25)A
(K16, K17, K23, G24, K25)A
10 µm

**Figure S10. Large-scale all-atom MD simulation of a SERF2 and rG4 complex. a** MD simulation of tetrameric TERRA10 (PDB: 2M18) and dimeric TERRA12 (PDB ID:2KBP) rG4 structures in 150 mM KCl, pH 7.4. The RNA, water molecules, $K^{+,}$ and $Cl^{-}$ atoms are shown in cartoon, red lines, purple and green spheres, respectively. The cartoon structure on right shows 10 superimposed structures of TERRA rG4 retrieved at 10 ns intervals. The TERRA rG4 structure at 0 ns and 100 ns are shown in dark orange and cyan, respectively. **b** Cartoon structure of SERF2-TERRA10 rG4 2:1 complex show N- and C-terminal charged residues in SERF2 interact with TERRA10 rG4. **c** 3D structure of TERRA10 rG4 complex with SERF2 before and after 0.5 μs MD simulation shows G-tetrad distortion in SERF2:TERRA10 rG4 1:1 (center) and 1:2 (right) complex. G-tetrads are indicated by red arrows and each TERRA10 rG4 unit in the tetrameric structure is represented with different colors. A G-quartet in TERRA10 rG4 is formed by guanines in i and i+6 as shown on the top. **d** Existence, and disappearance of hydrogen bonds in SERF2-TERRA10 rG4 1:1 (top) and 2:1 (bottom) complex as a function of adenosine uridine and guanine nucleotide concentration. The scale represents the total number of hydrogen bonds between SERF2 and TERRA10 rG4 in a given time frame. **e** Monitoring the change in TERRA12 rG4 (20 μM) parallel secondary structure (250-280 nm) by CD spectroscopy in the presence of an increasing amount of SERF2 as indicated. The full CD spectra are shown in the inset. CD spectra were collected at 25 °C for samples dissolved in 20 mM NaPi, 100 mM KCl, pH 7.4. **f** Phase transition of SERF2 and its mutants mixed with equimolar TERRA23 rG4s in a crowding condition (20 mM NaPi, pH 7.4, 100 mM KCl, 10% w/v PEG8000).

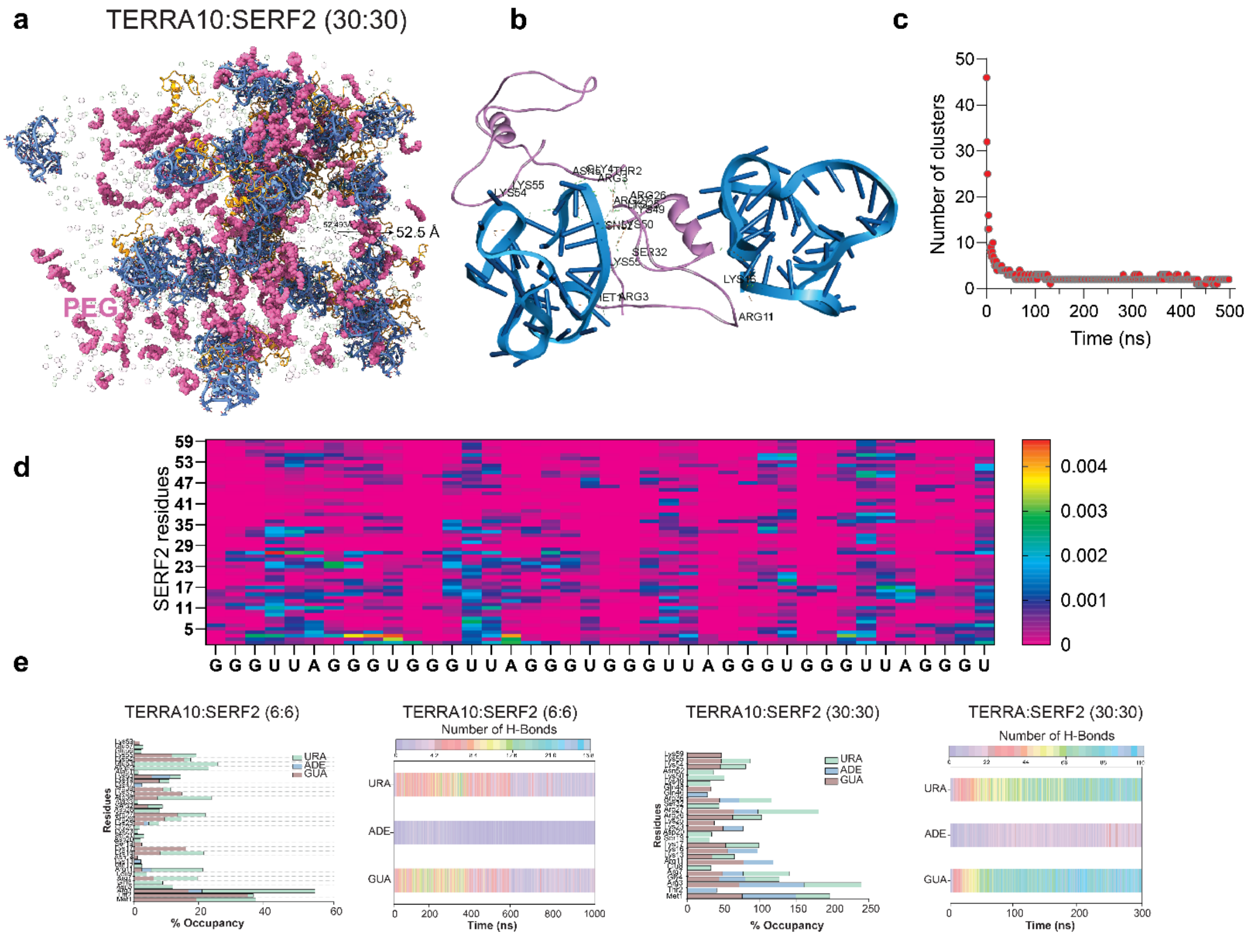


**Figure S11. Large-scale all-atom MD simulation of SERF2 and TERRA rG4 complex. a** MD snapshot retrieved at 0.5 μs for the large MD system comprising 30 SERF2 and 30 TERRA10 rG4 molecules, showing a hydrophilic core within the ring of size ~ 6 nm. The protein, rG4s, ions, and PEG crowders are shown in orange, blue, transparent spheres, and pink, respectively. Water molecules are not shown for clarity. **b** Cartoon structure of SERF2-TERRA10 rG4 2:1 complex show N- and C-terminal charged residues in SERF2 interact with TERRA10 rG4. **c** Cluster analysis using the MDAnalysis program shows that the randomly distributed 30 SERF2 and 30 TERRA10 rG4 molecules converge to generate a very few clusters during the 0.5 μs MD simulation. **d** Plot shows contact maps among 30 SERF2 and 30 TERRA10 rG4 molecules generated using MDanalysis program. **e** Interaction network between SERF2 and TERRA10 rG4 in large-scale all-atom MD simulations. SERF2 residue-specific interaction with TERRA10 rG4s presented as % occupancy, referring to the hydrogen bond occupancy as a function of simulation time in a small MD system containing 6 SERF2 and TERRA10 rG4 molecules, and in a large MD system consisting of 30 SERF2 and TERRA10 rG4 molecules. The number of hydrogen bonds in the corresponding small and large MD systems is shown on the right. The % occupancy over 100% means a single residue forming multiple bonds with the TERRA10 rG4 molecules.

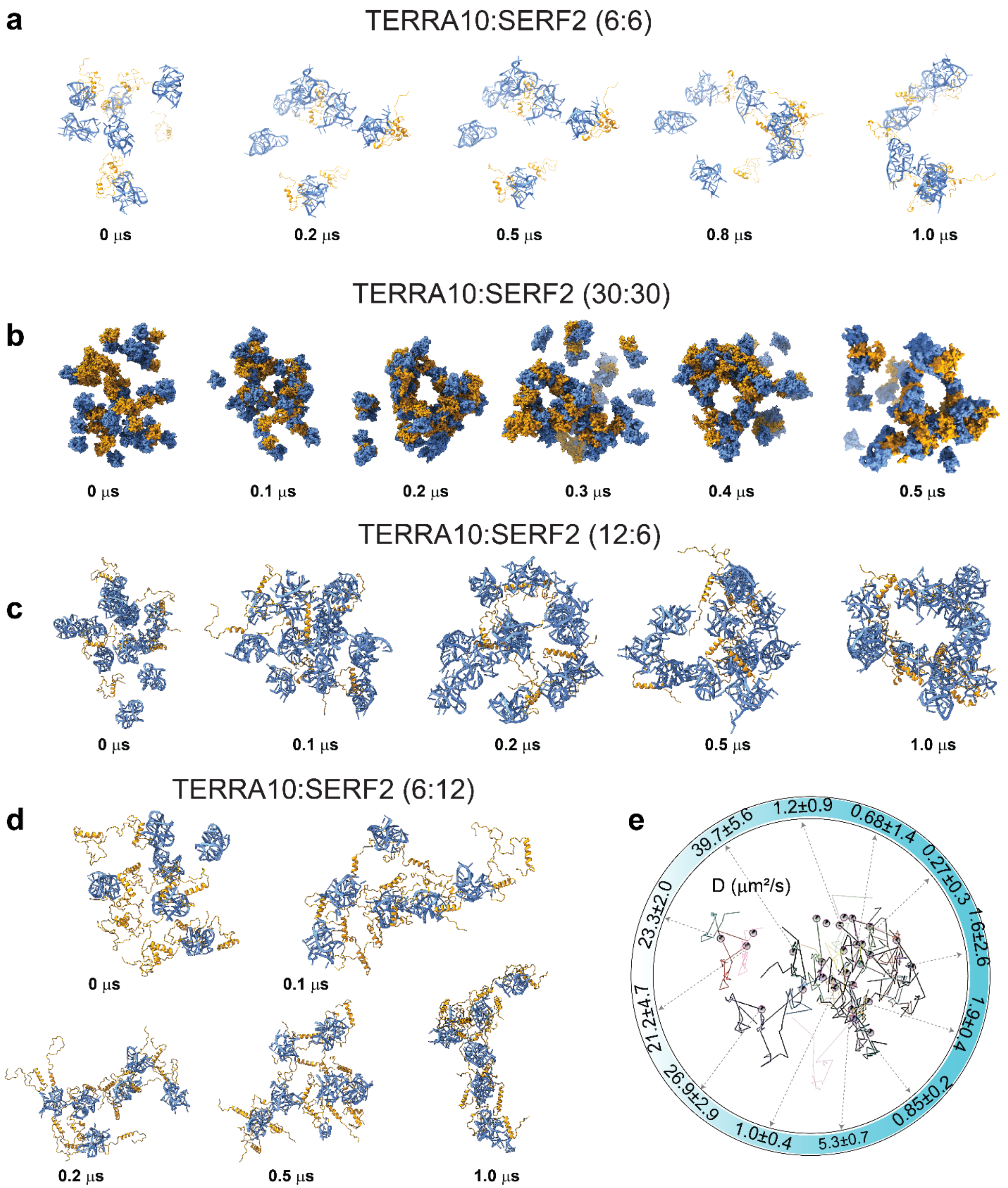


**Figure S12. Ring-like structure formation in SERF2–TERRA rG4 all-atom MD simulations. a-d** MD snapshots showing the formation of a ring-like structure retrieved at the indicated time-intervals in the large-scale MD simulation system comprising a small (**a**) or large (**b**) system of 1:1 protein to TERRA10 rG4 ratio, and a 2:1 (**c)** or 1:2 system (**d**). The SERF2 and TERRA rG4s molecules are shown in orange and blue, respectively. **e** Single-molecule tracking image constructed using the 0.5 µs all-atom MD trajectory in the 30 to 30 SERF2-TERRA rG4 all-atom MD simulation system. Each sphere represents the center of mass of individual SERF2 molecules, and their trajectories were retrieved at every 20 nanoseconds interval from the 0.5 μs MD simulation to construct the tracks using ImageJ. The diffusion constants of a set of representative SERF2 molecules in the lower-ordered dilute and droplet-like condensed phase, are indicated by arrows.

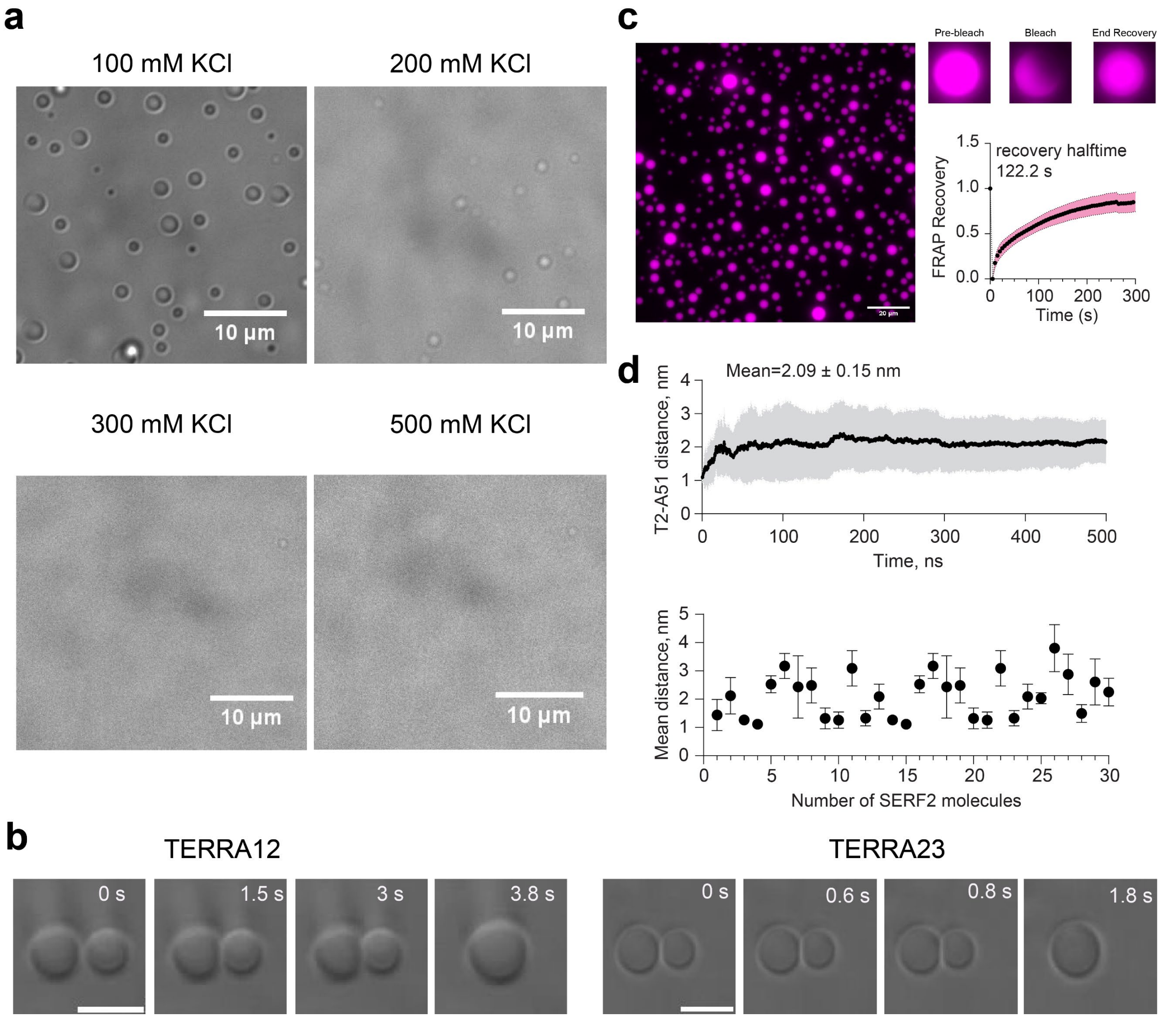


**Figure S13. Phase transition of SERF2 and TERRA rG4s. a** Effect of salt on SERF2 and TERRA12 rG4 phase transition. 50 μM of SERF2 was mixed with equimolar TERRA12 rG4 in 20 mM NaPi, pH 7.4 containing variable salt as indicated and 10% w/v PEG8000. **b** Measuring the SERF2 and TERRA12 or TERRA23 rG4 phase-separated droplet fusion time by a dual-trap optical tweezers. Scale bar is 5 μm. **c** Phase transition of 100 μM SERF2 containing 0.5% Cy5-labeled SERF2 (purple) mixed with equimolar TERRA12 rG4 dissolved in 20 mM NaPi, 100 mM KCl, pH 7.4. The droplet end recovery before and after FRAP is shown on the right. FRAP recovery plot of SERF2 (100 μM) in droplets containing equimolar TERRA12 rG4s (bottom right). The half-time was calculated by fitting the curve using a one-phase association model in GraphPad Prism. Standard deviations were calculated by analyzing 8 isolated droplets subjected to FRAP. **d** The plot on top shows the average end-to-end distance calculated from nitrogen atoms of T2 and A51 in each SERF2 molecule in the SERF2-TERRA10 rG4 0.5 μs all-atom MD system comprising 30 molecules of protein and rG4. Data are shown as mean values ± standard deviation. The plot on the bottom shows the mean distance ± standard error between T2 and A51 computed for individual SERF2 molecules.

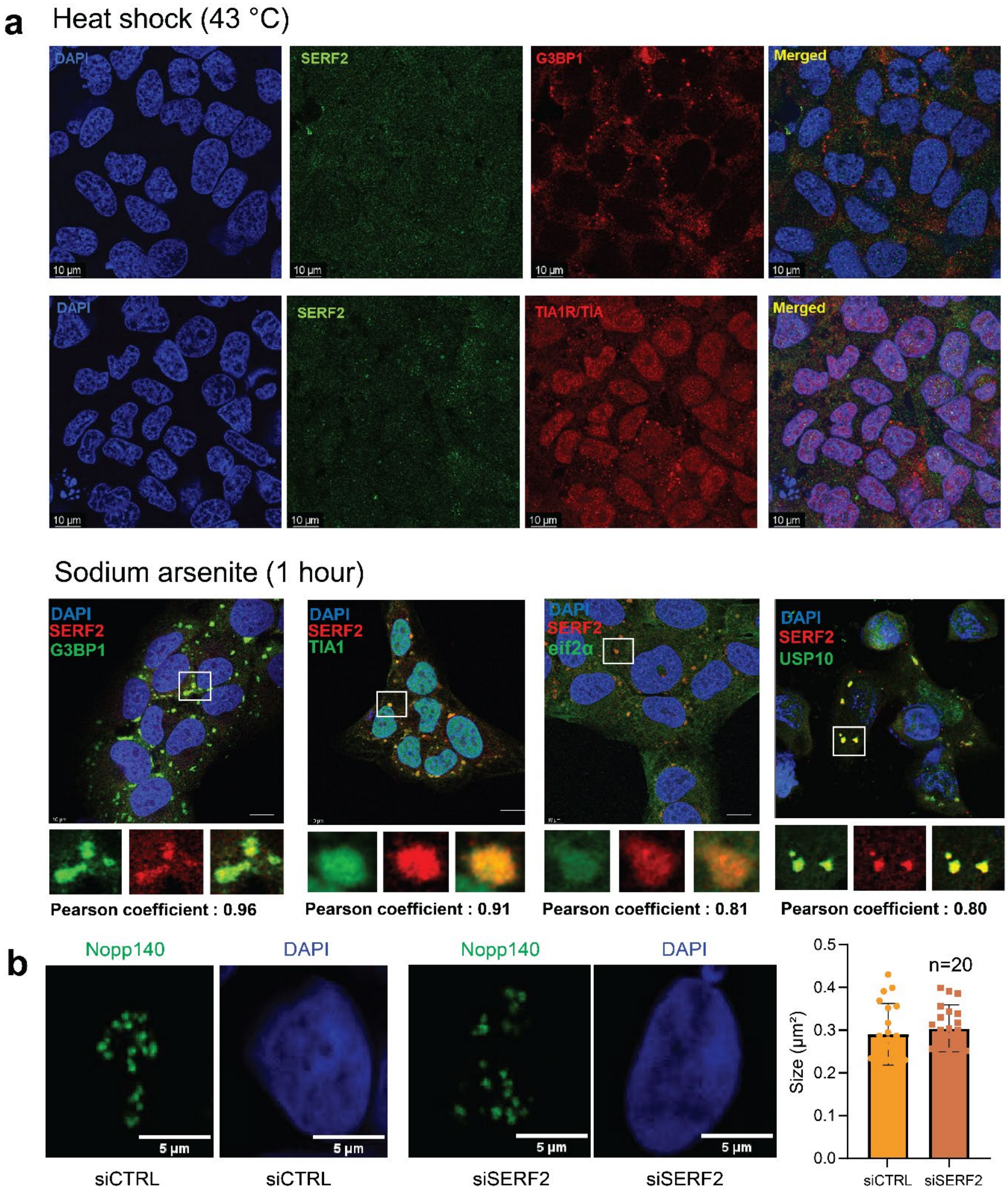


**Figure S14. SERF2 colocalized with different stress granule proteins. a** Immunofluorescence of SERF2, G3BP1 and TIA1 in fixed U2OS cells treated with heat shock (top panel), and SERF2, G3BP1, TIA1, eIF2α and USP10 treated with 0.5 mM sodium arsenite for 1 hour (bottom panel). Scale bar is 10 µm. **b** Comparing the size and shape distribution of nucleolus in cells treated with 0.4M sorbitol in siControl and siSERF2 conditions. Fixed cells were treated respectively with Nopp140 and DAPI to mark the nucleolus and nucleus. The plot on the right shows the size of twenty nucleoli (n=20) obtained from three replicates. Data are shown as mean values ± standard deviations.

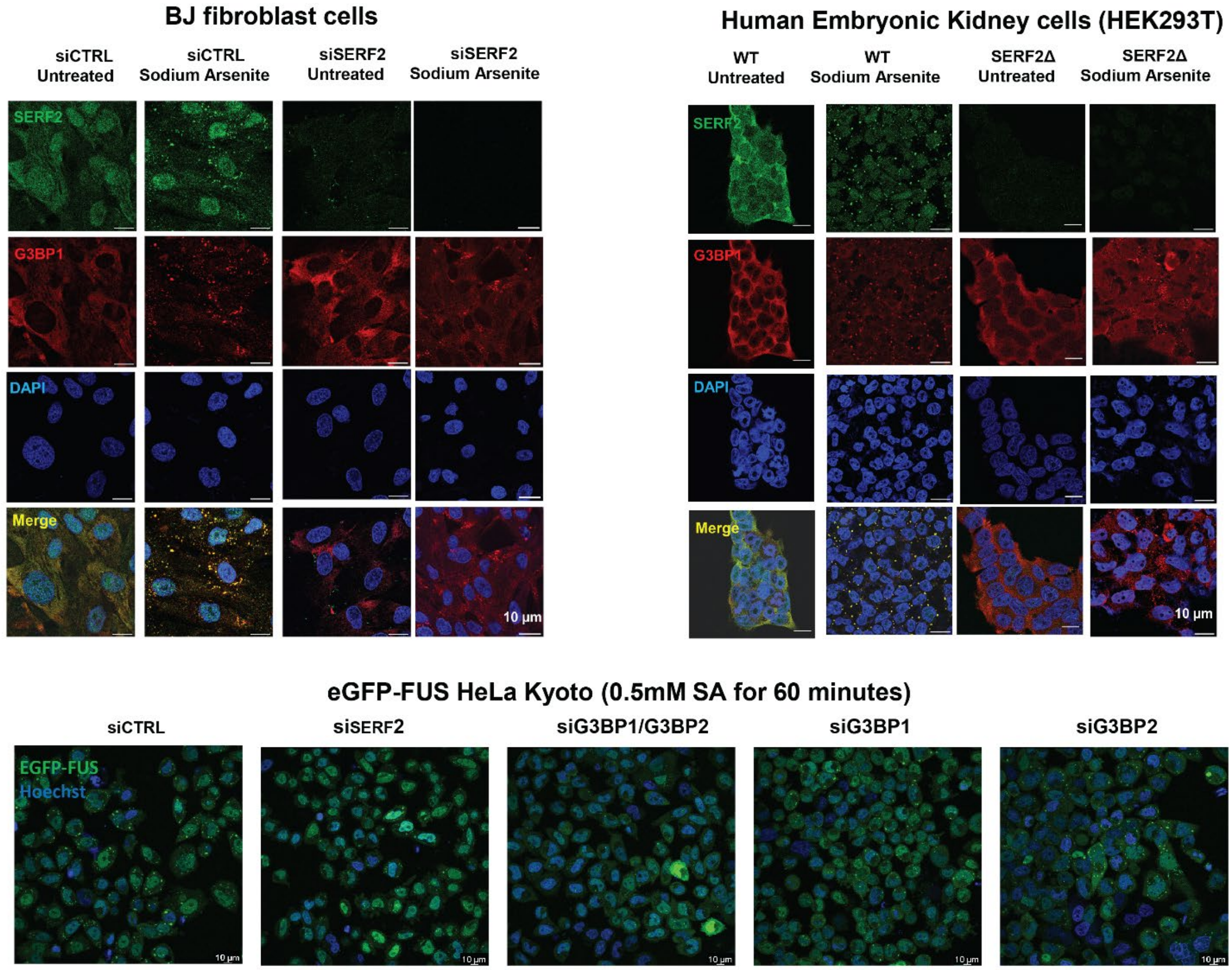


**Figure S15. SERF2 deletion or knockout affects stress granule assembly.** Immunofluorescence of SERF2 and G3BP1 in fixed BJ fibroblast cells with siCTRL or siSERF2 knockdown (top left), and SERF2 and G3BP1 in fixed HEK293T wild-type (WT) and SERF2 deletion cells (top right). Live cell imaging of EGFP-FUS HeLa Kyoto cells with siCTRL, siSERF2, siG3BP1/G3BP2, siG3BP1, or siG3BP2 knockdown (bottom). Cells were treated with 0.5 mM sodium arsenate for 30 minutes before imaging. Scale bar is 10 μm.

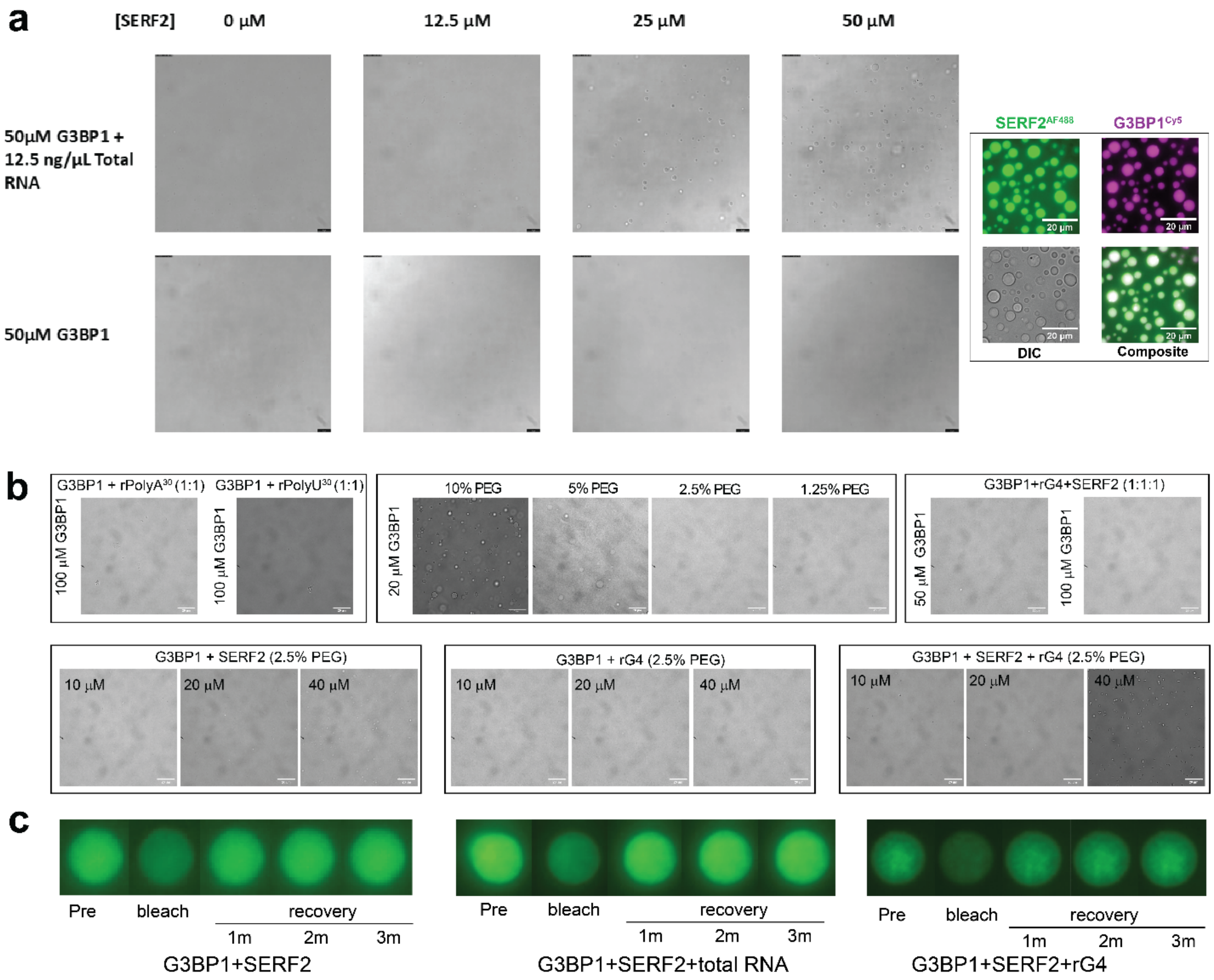


**Figure S16. The influence of SERF2 and RNA G-quadruplex or rG4 on G3BP1 phase transition. a** DIC images showing droplet formation in SERF2 mixed with 50 µM G3BP1 and 12.5 ng/µL of total RNA isolated from HeLa cells (top) at the indicated concentration. 50 µM G3BP1 (bottom) shows no droplets under the same conditions in the absence of SERF2 and total RNA. Scale bar for images on left panel is 10 µm. The right panel shows co-condensation of 50 µM G3BP1 mixed with 50 µM of SERF2 containing 0.5% Cy5-G3BP1 and AF488-SERF2. **b** DIC images showing the phase transition of G3BP1 at the indicated concentration in crowding or non-crowding environments mixed without or with RNA and SERF2 (top panel). DIC images show phase transitions of 20 µM G3BP1 suspended in 2.5% w/v PEG8000 mixed with an increasing concentration (10, 20, and 40 µM) of SERF2, TERRA23 rG4 and the mixture of SERF2 and rG4s (bottom panel). **c** Time-series FRAP recovery images of SERF2-AF488 in different condensates containing G3BP1, total RNA, or TERRA23 rG4s as indicated. The FRAP recovery plots are shown in Figure 7e.
